# *Aedes aegypti* and *Aedes albopictus* abundance, landscape coverage and spectral indices effects in a subtropical city of Argentina

**DOI:** 10.1101/2022.01.11.475665

**Authors:** Mía Elisa Martín, Ana Carolina Alonso, Janinna Faraone, Marina Stein, Elizabet Lilia Estallo

**Affiliations:** Instituto de Investigaciones Biológicas y Tecnológicas, Universidad Nacional de Córdoba. Av. Vélez Sarsfield 1611, CP X5016GCA. Córdoba, Córdoba, Argentina; Consejo Nacional de Investigaciones Científicas y Técnicas, Argentina; Instituto de Medicina Regional, Universidad Nacional del Nordeste. Avda. Las Heras 727, CP 3500. Resistencia, Chaco, Argentina

**Keywords:** landscape, *Aedes aegypti*, *Aedes albopictus*, public health, urban ecology, urban environment, dengue, Eldorado city

## Abstract

The presence, abundance and distribution of *Aedes* (*Stegomyia*) *aegypti* (Linnaeus 1762) and *Aedes* (*Stegomyia*) *albopictus* (Skuse 1894) could be conditioned by different data obtained from satellite remote sensors. In this paper, we aim to estimate the effect of landscape coverage and spectral indices on the abundance of *Ae. aegypti* and *Ae. albopictus* from the use of satellite remote sensors in Eldorado, Misiones, Argentina. Larvae of *Aedes aegypti* and *Ae. albopictus* were collected monthly from June 2016 to April 2018, in four outdoor environments: tire repair shops, cemeteries, family dwellings, and an urban natural park. The proportion of each land cover class was determined by Sentinel-2 image classification. Furthermore spectral indices were calculated. Generalized Linear Mixed Models were developed to analyze the possible effects of landscape coverage and vegetation indices on the abundance of mosquitoes. The model’s results showed the abundance of *Ae. aegypti* was better modeled by the minimum values of the NDVI index, the maximum values of the NDBI index and the interaction between both variables. In contrast, the abundance of *Ae. albopictus* has to be better explained by the model that includes the variables bare soil, low vegetation and the interaction between both variables.

## Introduction

In the world, the most important mosquito species in terms of disease transmission to humans are: *Aedes* (*Stegomyia*) *aegypti* (Linnaeus 1762) and *Aedes* (*Stegomyia*) *albopictus* (Skuse 1894). The arboviruses transmitted by these mosquitoes cause some of the most important diseases in the world (dengue, yellow fever, Zika, chikungunya and others), representing one of the greatest concerns for public health due to the great global interconnection mainly due to human population migrations, tourism, the growth of the transport of food and products, environmental changes related to urbanization, deforestation and climate change, among others (Juliano & Lounibos, 2005; Rúa-Uribe *et al*., 2012). These mosquito species are present in urban, suburban, and rural settlements in tropical, subtropical and temperate regions due to their ability to inhabit both natural (e.g., tree holes) and artificial (e.g., manholes, water storage containers, flower pots, used tires) breeding sites (Hawley, 1998; Vezzani & Carbajo, 2008). In particular, the distribution of *Ae. aegypti* include tropical, subtropical and temperate regions of the world, where it is considered an anthropophilic mosquito and is present mainly inside homes in urban areas. *Aedes albopictus* is distributed in the tropics worldwide, but also in temperate regions in the northern hemisphere, and is associated with the peri-domicile of suburban and rural environments (Lima-Camara *et al*., 2006, Robert *et al*., 2020).

In Argentina, since the first record in the country during the first half of the 20th century, *Ae. aegypti* was present in several provinces of the country. Currently, it is present in 19 provinces: Buenos Aires, Catamarca, Chaco, Córdoba, Corrientes, Entre Ríos, Formosa, Jujuy, La Pampa, La Rioja, Mendoza, Misiones, Neuquén, Salta, San Juan, San Luis, Santa Fe, Santiago del Estero and Tucumán (Grech *et al*., 2012; Rossi, 2015; Páez *et al*., 2016). The first record of *Ae. albopictus* in Argentina dates from 1998 when it was found in the cities of San Antonio and Eldorado in Misiones province (Rossi *et al*., 1999; Schweigmann *et al*., 2004). For 20 years, it had only been detected in three other cities in Misiones (Puerto Iguazú, Comandante Andresito, and Colonia Aurora) (Vezzani & Carbajo, 2008; Lizuain *et al*., 2019). At present, it has been found for the first time in Corrientes province in 2019, 200 km to the south from its previous records, representing the southernmost distribution in South America (Goenaga *et al*., 2020).

The presence, abundance and distribution of *Ae. aegypti* and *Ae. albopictus* could be conditioned by the landscape coverage from the differences presented in the biology, ecology and development of these vectors (Mudele & Gamba, 2019; Mudele *et al*., 2021). Changes in environmental conditions as a result of urbanization have been related (directly or indirectly) to the availability of breeding sites, and the modification in the abundance, richness, development and survival of adult mosquitoes (Baldacchino *et al*., 2017; Benitez *et al*., 2020). Different data obtained from satellite remote sensors have been used to indicate and identify favorable breeding sites for mosquitoes (Hassan *et al*., 2013). Some studies have linked mosquito populations to remotely detected land cover features. Vanwambeke *et al*. (2007) found a high probability of finding larvae of *Ae. albopictus* in the peri-urban. It has also been related to the presence of mixed areas of urbanization and vegetation (Manica *et al*., 2016). While the abundance and distribution of *Ae. aegypti* has been related to a greater extent, with variables related to urbanization, such as the presence of buildings (Sallam *et al*., 2017; Benitez *et al*., 2019).

On the other hand, vegetation is one of the most important and frequently described environmental characteristics in the spatial analysis of these species, being repeatedly used in research based on the calculation of satellite spectral indices (Heinisch *et al*., 2019). Numerous indices can be obtained from algorithms applied on the original remote sensor bands, two of these are potentially indicative of the presence of mosquito breeding sites due to the dependence of the immature stages on the aquatic habitat (Vanwambeke *et al*., 2007). The Normalized Difference Vegetation Index (NDVI) is the spectral vegetation index most used in spatial and temporal studies (Estallo *et al*., 2018; Benitez *et al*., 2019). Along with this, the Normalized Difference Water Index (NDWI) have been widely used in mosquito studies for many years (Pope *et al*., 1994; Mudele & Gamba, 2019), as well as applied in the study of vector-borne diseases (Estallo *et al*., 2012).

For Argentina, although the knowledge about the biology of *Ae. aegypti* is well documented (Carbajo *et al*., 2006; Estallo *et al*., 2018; Benitez *et al*., 2019), there is very little work on *Ae. albopictus* since its detection in 1998 (Schweigmann *et al*., 2004; Lizuain *et al*., 2019; Faraone *et al*., 2021). In this context and due to the absence of vaccines for most of the viruses transmitted by these two species, vector management and control is the main current tool to prevent their spread. Therefore, the aim of this study was to estimate the effect of landscape coverage and spectral indices on the abundance of *Ae. aegypti* and *Ae. albopictus* from the use of satellite remote sensors in Eldorado, Misiones, Argentina.

## Materials and Methods

### Study site

Eldorado city (Fig. 1) is located in the northwest of Misiones province, within the Neotropical region (26° 24′ S, 54° 38′ W). The phytogeographical region is Paraná province. The area characterized by the presence of three arboreal strata, with lianas, epiphytes and hemiepiphytes and an undergrowth of ferns and herbaceous and shrubby phanerophytes, including bamboos (Oyarzabal *et al*., 2018). The climate is subtropical, hot and humid, without a marked dry season. The mean annual temperature is 22 °C, with a maximum temperature of 38.5 °C (January) and a minimum of 5.4 °C (July); the mean annual rainfall is 2020 mm (Silva *et al*., 2008).

**Fig. 1.**
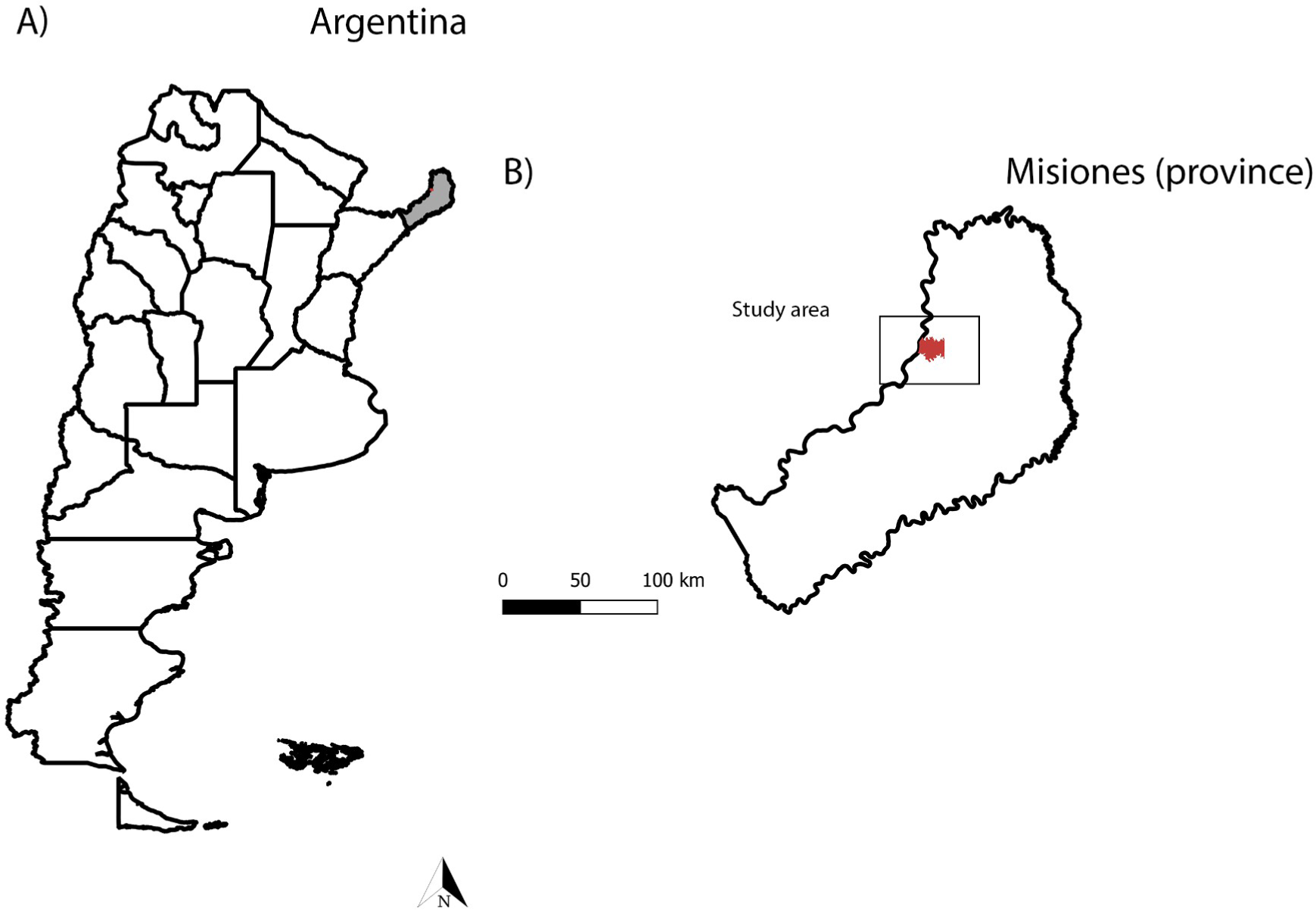
(A and B) Geographic location of the study area in Misiones, Argentina.

Eldorado is the third-largest city in the province with a population of 100,000 inhabitants and a surface of 215 km^2^ where 14% corresponds to rural areas, 30.6% to natural forests and 55.4% to other uses (Molinatti *et al*., 2010). The city expands on both sides along the National Route N° 12. The main economic activities of the region are forestry (sawmills, pulp and paper industry) and agriculture, oriented to industrial crops production of (yerba mate, tea, tobacco and citrus).

### Entomological sampling

Larvae of *Aedes aegypti* and *Ae. albopictus* were collected monthly from June 2016 to April 2018, in four outdoor environments: tire repair shops, cemeteries, family dwellings, and an urban natural park (Parque Schwelm) (Fig. 2). Sampling sites with larval presence of both species were georeferenced using the Global Position System (GPS-Garmin eTREX 10). The number of monthly samples was N = 60, distributed as follows: 20 natural habitats; 20 artificial habitats of cemeteries, 10 of repair shops and 10 of houses. The homes were visited according to the provisions of the Environmental Sanitation Direction of the Municipality of Eldorado, where each month different neighborhoods were visited. The larvae were transferred to the laboratory of the Institute of Regional Medicine for their breeding (larvae of instar I, II and III), conservation and determination. For morphological identification of the specimens (fourth instar larvae), dichotomous keys (Darsie 1985; Consoli & de Oliveira 1994) were used.

**Fig. 2.**
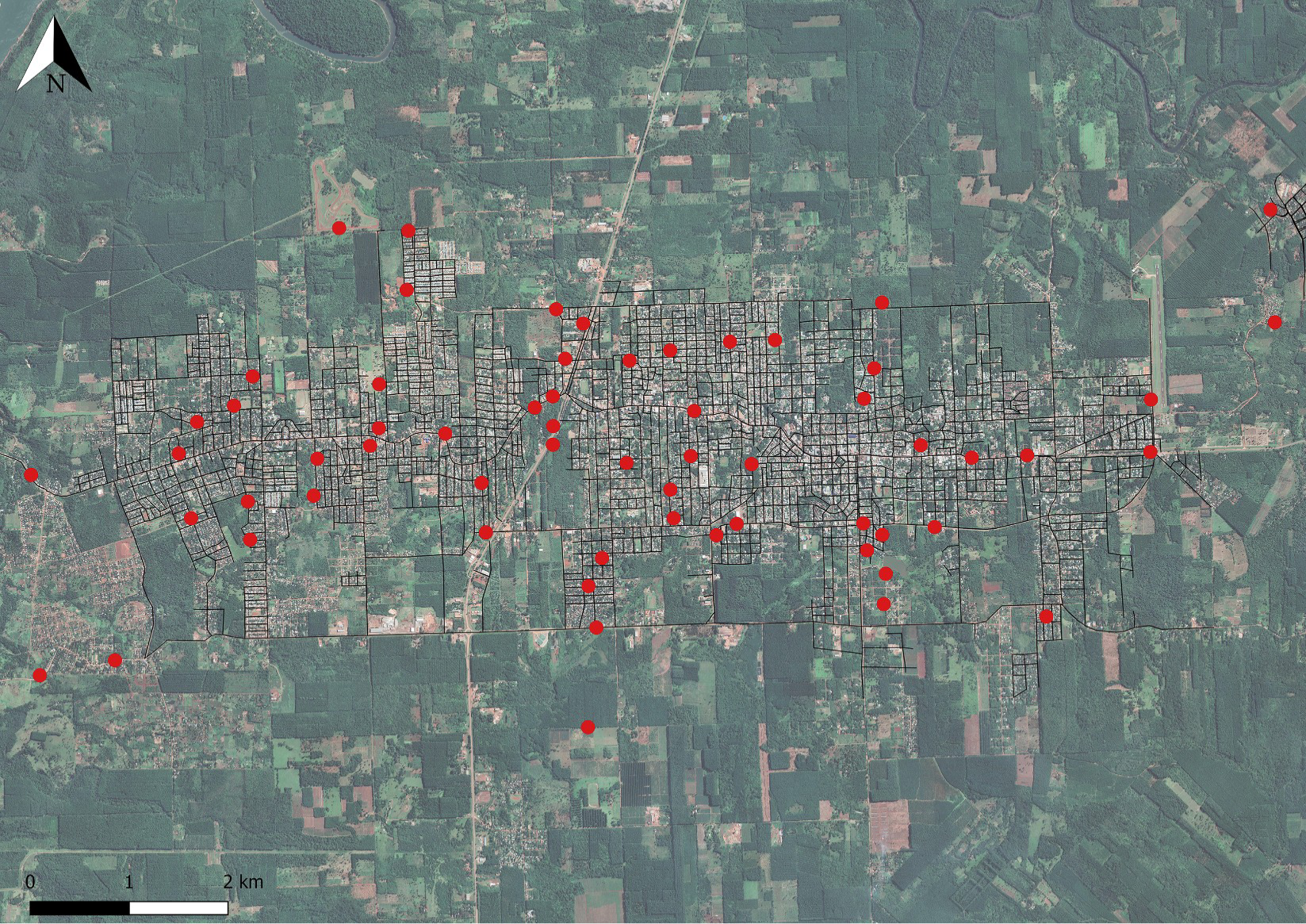
Distribution of sampling sites in Eldorado, Misiones, Argentina.

### Remote sensing data

In order to estimate the different landscape coverage in the city, images from the Sentinel-2 satellite were used. Five images from the satellite were used, which were downloaded from the Land Viewer website (https://eos.com/landviewer/). The satellite images correspond to the succession of stations from the three years of sampling and were selected according to the availability of images on the website and the absence of clouds over the area of interest.

#### Spectral indices

On each satellite image, spectral indices were calculated: Normalized Difference Vegetation Index (NDVI), Normalized Difference Water Index (NDWI) and Normalized Difference Built-up Index (NDBI). The NDVI reflects the contrast of vegetation reflectivity between the spectral regions of Red (R) and Near Infrared (NIR) reflectance (Eq.1). This index can be associated with the vegetation cover, in terms of abundance and vigor, since it is strongly related to the photosynthetic activity of the vegetation, allowing to identify the presence of vegetation on the surface and characterize its spatial distribution. The values vary from −1 to +1, where high values correspond to areas with vigorous vegetation, negative values are associated with covers such as water and values close to zero correspond to bare soil (Chuvieco Salinero, 2008). On the other hand, the NDWI is an index that takes into account the water content present in the mesophyll of the leaves and indirectly measures precipitation and soil humidity (Estallo *et al*., 2012). It varies between −1 and +1, depending on the water content of the leaves, but also on the type of vegetation and cover. It is based on the contrast between the reflectances of Short-wave Infrared (SWIR) and NIR wavelengths (Eq.2) (Gao 1996). The NDBI is an index that highlights urban areas, where there is typically a higher reflectance in the SWIR region, compared to the NIR region (Zha *et al*., 2003). Positive NDBI values indicate built-up areas and those close to 0 indicate vegetation, while negative values represent bodies of water (Ranagalage *et al*., 2017).

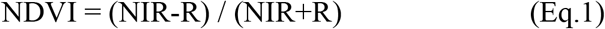

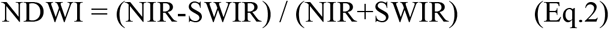

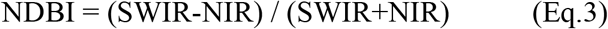

#### Land cover classification

To determine landscape coverage in Eldorado, supervised classification (Minimum Distance to Mean) was performed using QGIS 3.4.15 software (https://www.qgis.org/). Five land cover classes were obtained: water (rivers, lakes, artificial bodies of water), bare soil (soil without any vegetation cover, unpaved streets), urban areas (buildings, paved streets and roads), low vegetation (herbs and grasses) and high vegetation (trees and shrubs). The accuracy of the classification was measured by a confusion matrix and the value of the Kappa’s coefficient, where values close to 1 indicate greater accuracy of the classification method. The areas for verification were determined from the visualization of images published in Google Earth ©. A total of 100 control points were defined by landscape coverage following the criteria recommended by Chuvieco Salinero (2008). Regarding the classification of Sentinel-2 images, the global precision of the classifications ranged from 91% to 99.6%, with Kappa’s coefficients from 0.887 to 0.995. One of the final classified images can be seen in Figure 3.

**Fig. 3.**
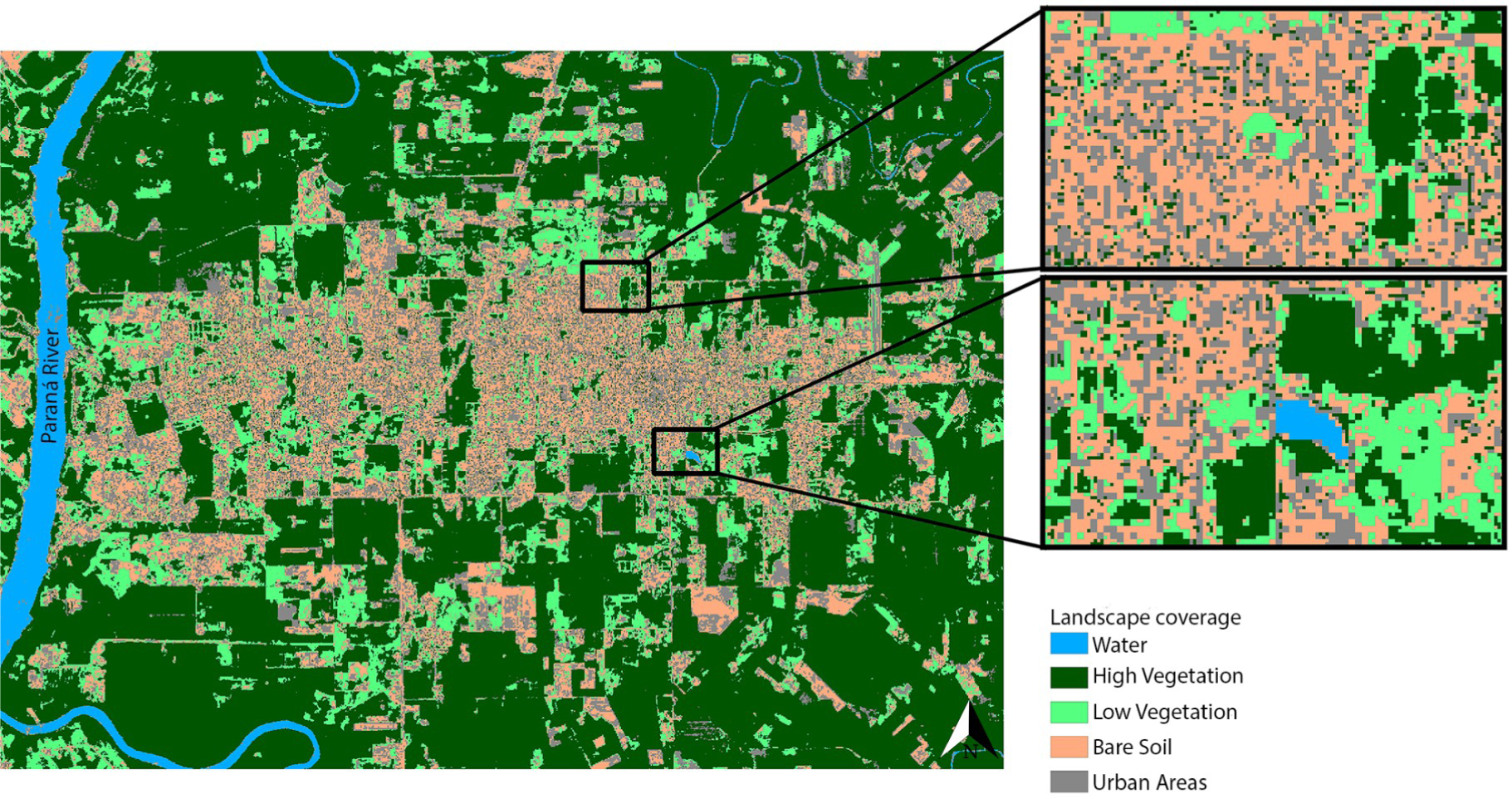
Supervised classified image for Eldorado from November 12, 2016.

#### Buffer areas

Around each sampling site, circular influence areas of 100m were generated, avoiding the overlapping of the areas and taking into account the biology of the vector. Once these areas were constituted in each classified image, the proportions of each class of landscape coverage were extracted, as well as the mean, minimum and maximum values of NDVI, NDWI and NDBI.

#### Data analysis

To analyze the possible effects of landscape coverage and vegetation indices on the abundance of larvae, generalized linear mixed models (GLMM) were constructed for each species separately with a Negative Binomial distribution. To control for over-scattering, a logarithmic link function was used (Zuur *et al*., 2009). In our analyzes, the response variable used was the number of larvae collected at each site per month. The sites were incorporated as a random effect to include spatial dependence. The explanatory variables used are shown in Table 1. Water coverage was not incorporated into the models because it was not found in any buffer area.

**Table 1.** Explanatory variables used to explain the variation in the abundances of *Ae. aegypti* and *Ae. albopictus* in Eldorado, Misiones.

| Variable | Description |
| --- | --- |
| highV | Proportion of high vegetation cover extracted from a 100m buffer around each sampling site |
| lowV | Proportion of low vegetation cover extracted from a 100m buffer around each sampling site |
| soil | Proportion of bare soil cover extracted from a 100m buffer around each sampling site |
| urban | Proportion of urban areas cover extracted from a 100m buffer around each sampling site |
| ndvi | Mean value of NDVI extracted from a 100m buffer around each sampling site |
| ndvimin | Minimum value of NDVI extracted from a 100m buffer around each sampling site |
| ndvimax | Maximum value of NDVI extracted from a 100m buffer around each sampling site |
| ndwi | Mean value of NDWI extracted from a 100m buffer around each sampling site |
| ndwimin | Minimum value of NDWI extracted from a 100m buffer around each sampling site |
| ndwimax | Maximum value of NDWI extracted from a 100m buffer around each sampling site |
| ndbi | Mean value of NDBI extracted from a 100m buffer around each sampling site |
| ndbimin | Minimum value of NDBI extracted from a 100m buffer around each sampling site |
| ndbimax | Maximum value of NDBI extracted from a 100m buffer around each sampling site |

First, data exploration was implemented following the protocol described in Zuur *et al*. (2010). The explanatory variables were standardized to balance their weight and also to avoid introducing errors in the model produced by the different measurement units of each variable. Then, a Spearman’s test was performed to analyze the correlation of the explanatory variables.

The models were built using a manual step-by-step forward procedure. We began by evaluating the significance of each response variable from univariate GLMM. The variables that were significant for each species were in turn used as starting points in the different branches of the modeling. Subsequent variables were added one at a time as long as they did not have a correlation coefficient >0.7 with some variables already included. Interactions between them were also tested. In each step, the significance of each addition was evaluated with a significant reduction (2 points) in the Akaike Information Criterion corrected for low sample sizes (AICc) (Zuur *et al*., 2009). The GLMMs were classified according to the AICc and the model with the lowest value was selected as the best model. The multicollinearity between variables was evaluated in the final models using the Variance Inflation Factor, considering a threshold value equal to 5. Finally, the ggResidpanel package was used to verify the normality of the residual distribution and evaluate the residual plot.

The free software R, version 4.0.3 (https://www.r-project.org/) and the packages lme4 (*glmer*.*nb* function), MuMin (*model*.*sel* function) and car (*vif* function) were used to perform the statistical analyzes.

## Results

A total of 23,658 mosquitoes of the species under study were collected during the entire sampling period. Of that total, *Ae. aegypti* presented a relative abundance of 86.70% (n = 20,511), while *Ae. albopictus* of 13.30% (n = 3147).

Based on the exploratory analysis of the variables and considering those with statistical significance in the univariate GLMMs, 5 model branches were constructed for *Ae. aegypti* and 1 branch for *Ae. albopictus*. For the first species, the univariate GLMMs of: highV, soil, ndvimin, ndbi and ndbimax were started, and after considering the correlations between the independent variables, 66 models were made that evaluated the addition of more variables and interactions. In contrast, for *Ae. albopictus* GLMMs were modeled from the variable: soil, making 14 models (see Tables A-G in Supporting Information). In Table 2, the selected models within each branch are displayed from the comparison of the goodness of fit indicators (AICc) for the species under study.

**Table 2.**
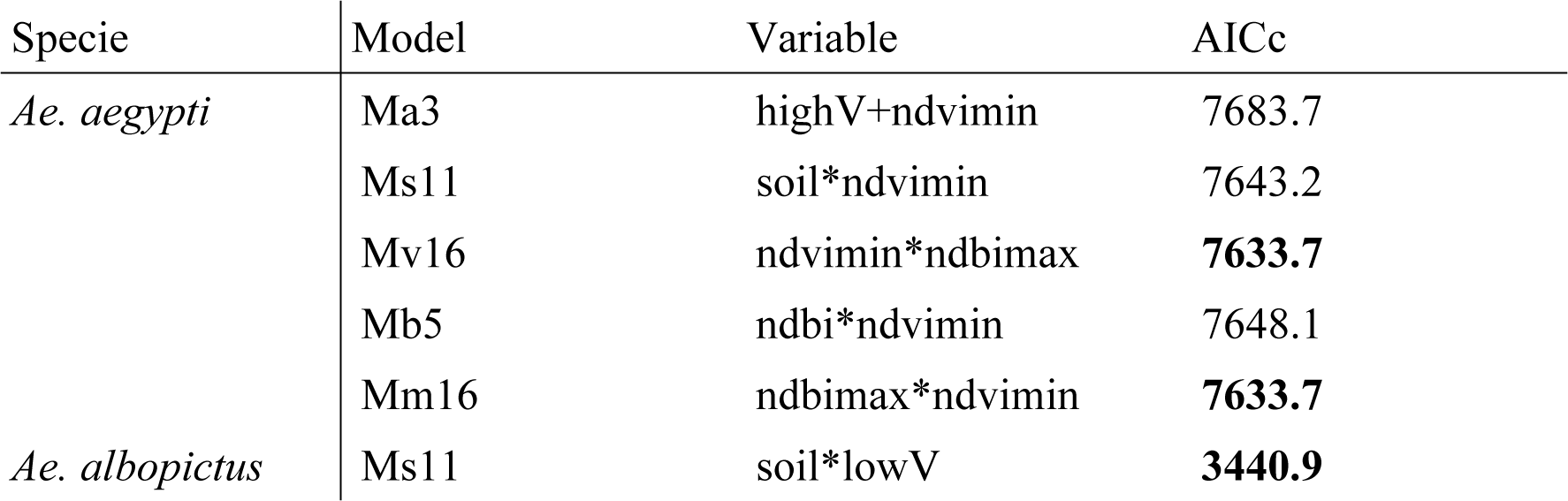
GLMM selected for *Ae. aegypti* and *Ae. albopictus*.

| Specie | Model | Variable | AICc |
| --- | --- | --- | --- |
| *Ae. aegypti* | Ma3 | highV+ndvimin | 7683.7 |
|  | Ms11 | soil*ndvimin | 7643.2 |
|  | Mv16 | ndvimin*ndbimax | **7633.7** |
|  | Mb5 | ndbi*ndvimin | 7648.1 |
|  | Mm16 | ndbimax*ndvimin | **7633.7** |
| *Ae. albopictus* | Ms11 | soil*lowV | **3440.9** |

The GLMM results showed that the larvae abundance of *Ae. aegypti* was better modeled by the minimum values of the NDVI index, the maximum values of the NDBI index and the interaction between both variables (Table 3). In contrast, the abundance of *Ae. albopictus* has to be better explained by the model that includes the variables soil, lowV and the interaction between both variables (Table 4). The other GLMM with the same AICc (soil*ndbimax) was not selected for presenting a vif >5 in the interaction between the variables.

**Table 3.** Coefficients of the final GLMM selected for *Ae. aegypti*.

| Variable | Estimate | Std. Error | Z value | Pr(>\|z\|) |
| --- | --- | --- | --- | --- |
| Intercept | 5.18544 | 0.01475 | 351.6 | <2e-16** |
| ndvimin | -14.48629 | 0.01481 | -977.8 | <2e-16** |
| ndbimax | -6.94077 | 0.01481 | -468.6 | <2e-16** |
| ndvimin*ndbimax | 17.40568 | 0.01482 | 1174.7 | <2e-16** |

An asterisk means $p < 0.05$ , two asterisks mean $p < 0.01$ .

**Table 4.**
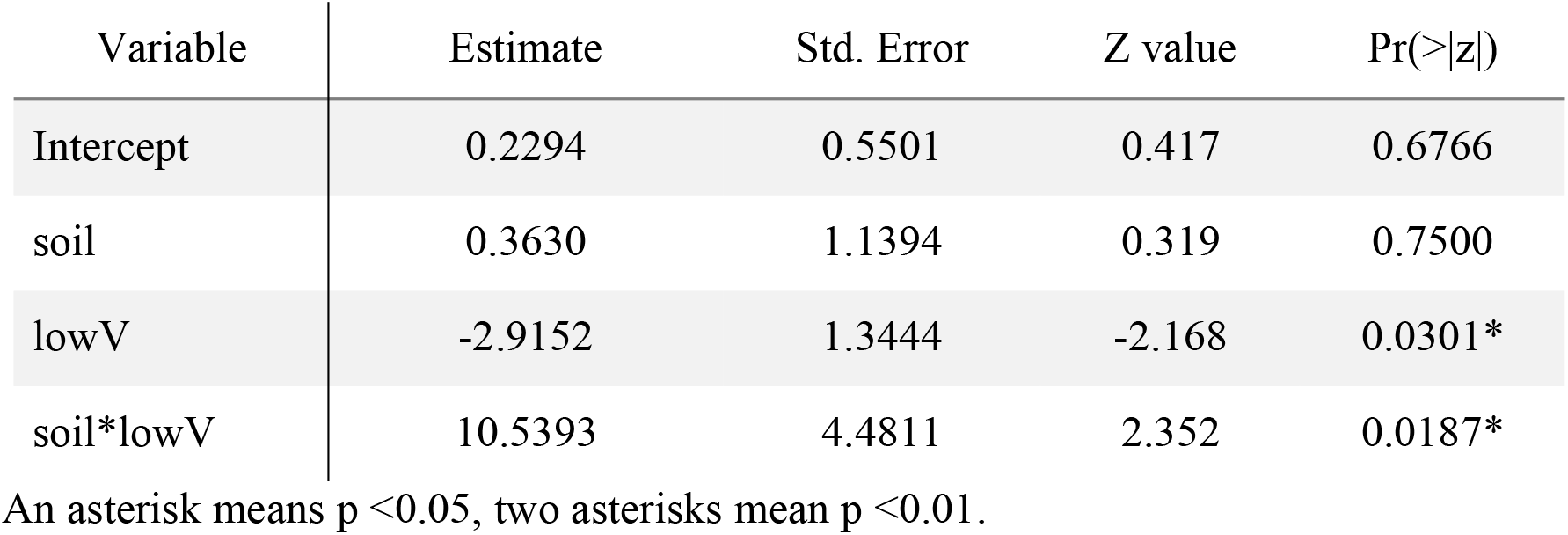
Coefficients of the final GLMM selected for *Ae. albopictus*.

| Variable | Estimate | Std. Error | Z value | Pr(>\|z\|) |
| --- | --- | --- | --- | --- |
| Intercept | 0.2294 | 0.5501 | 0.417 | 0.6766 |
| soil | 0.3630 | 1.1394 | 0.319 | 0.7500 |
| lowV | -2.9152 | 1.3444 | -2.168 | 0.0301* |
| soil*lowV | 10.5393 | 4.4811 | 2.352 | 0.0187* |

An asterisk means $p < 0.05$ , two asterisks mean $p < 0.01$ .

## Discussion

The present study allowed us to identify the effect of landscape covers and vegetation indices on the spatio-temporal larvae abundance of *Ae. aegypti* and *Ae. albopictus* from the use of Sentinel-2 images in a subtropical city of Misiones, Argentina.

The global distribution of ecological rivals, *Ae. aegypti* and *Ae. albopictus*, have changed in recent decades due to differences in their abilities to compete with each other (Bennett *et al*., 2021). Generally, *Ae. aegypti* is highly adapted to the domestic environment, and therefore abundance is positively correlated with increasing urbanization (Higa, 2011). In this study, a negative association was found between the abundance of *Ae. aegypti* and NDVI minimun values and NDBI maximun values. In accordance with Bennett *et al*., (2021) who found a negative association with lower NDVI values for both species in Panamá.

Urban areas provide this mosquito with food, shelter, reproduction and oviposition sites (Flaibani *et al*., 2020). Previous studies in the United States, Costa Rica, Puerto Rico, Brazil and Argentina, have related the abundance of the species with urban areas, buildings and high housing density (Carbajo *et al*., 2006; Vezzani & Carbajo, 2008; Fuller *et al*., 2010; Little *et al*., 2011; Montagner *et al*., 2018; Benitez *et al*., 2019, Heinisch *et al*., 2019). In turn, Chaves *et al*., (2021) in Costa Rica found a negative association between vegetation index (measured through the Enhanced Vegetation Index-EVI-) and the abundance of *Ae. aegypti*, while Samson *et al*., (2015) found that urban areas identified by Urban Index were found to be important in predicting distribution of the species and that the results of their models show a high probability for *Ae. aegypti* in and around urban areas. In accordance with our findings about the negative association of *Ae. aegypti* with the maximum values of NDBI, a spatial study carried out in Buenos Aires city, Argentina found that the proliferation of mosquitoes *Ae. aegypti* was highest in medium urbanization levels (not densely built on the suburban areas) (Carbajo *et al*., 2006). Due to the different population densities of both cities, we expect that the maximum values of NDBI in Eldorado (57,323 inhabitants) will be related to the mean values of NDBI in Buenos Aires (12,801,364 inhabitants). In cities with a high degree of urbanization and high population density, the peripheral area is the most conducive to the reproductive activity of the vector since urbanized areas of the city offer few spaces with vegetation (for food and shelter), few breeding sites and reduce the connectivity between patches of habitat that are more favorable (Carbajo *et al*., 2006, Benitez *et al*., 2019)

Our study found a positive association between the abundance of *Ae. aegypti* and the interaction between both indices in accordance with previous studies for Costa Rica (Troyo *et al*., 2009) and temperate Argentina area (Benitez *et al*., 2019) where moderately built-up residential areas with moderate tree cover likely contain a relatively high number of positive habitats for this species, therefore heterogeneity in urban areas can be linked to the distribution of this species.

On the other hand, the distribution of *Ae. albopictus* is associated with vegetation in rural, suburban and urban areas and its abundance is negatively affected by urbanization. This difference in distribution along the urban-rural gradient is associated with behavior related to blood feeding, host preference, and preference for vegetation, offering ideal conditions for resting and egg laying (Heinisch *et al*., 2019; Higa, 2011; Manica *et al*., 2016). We observed a negative association between the abundance of the species and low vegetation coverage, and a positive association between the interaction of soil and low vegetation. In this work, the land cover class soil has been related around the sampling sites with sandy streets (unpaved road) more characteristic of suburban areas (see Fig. A-C in Supporting Information). Our results are according to Myer *et al*. (2019), who found an important relationship between the abundance of *Ae. albopictus* and grass cover (negative) and the interaction between impervious and grass cover (positive).

In agreement with Rey *et al*. (2006), Honorio *et al*. (2009) and Cianci *et al*. (2015) low vegetation coverage that includes grasses was negatively associated with the abundance of *Ae. albopictus* larvae, indicating that open areas are less attractive for this mosquito species. In Porto Alegre, Brazil, *Ae. albopictus* was dominant in urban areas with vegetation, relating its adaptation to transition zones between urban and non-urban/natural habitats (Montagner *et al*., 2018). According to Forattini (2002), the adaptation of the species to transition zones results from being able to use larval habitats or breeding sites and sources of blood food from both environments. Likewise, in Florida, United States, Rey *et al*. (2006) found a positive relationship between the abundance of immature *Ae. albopictus* and land covers: ground vegetation, unpaved road and bare ground.

This is the first work carried out in the country to relate the abundance of *Ae. albopictus* with products derived from remote sensors, and the results obtained provide important knowledge about the biology of this species in Argentina. The Pan American Health Organization (PAHO, 2016) has recommended the following in areas of recent infestation by *Ae. albopictus* the immediate responsibility to contain and control it if possible, to prevent further spread. For this, knowledge is required on numerous aspects of the ecology of the species, areas of distribution, periods of greater activity, among others that generate baselines to understand the dynamics of pathogen transmission and therefore implement effective programs of control.

## Acknowledgements

We thank Mr. Carlos Paredes and the technicians from the Environmental Sanitation Direction of the Municipality of Eldorado for their technical support in collecting specimens.

## Author Contributions

Mia E. Martin: Formal analysis, Writing - original draft, Writing - review & editing. Ana C. Alonso: Investigation, Methodology, Writing - original draft, Writing - review & editing. Janinna Faraone: Investigation, Methodology, Writing - original draft, Writing - review & editing. Marina Stein: Conceptualization, Funding acquisition, Methodology, Investigation, Resources, Supervision, Writing - original draft, Writing - review & editing. Elizabet L. Estallo: Formal analysis, Methodology, Writing - original draft, Writing - review & editing.

