## Supporting Information for "*Aedes aegypti* and *Aedes albopictus* abundance, landscape coverage and spectral indices effects in a subtropical city of Argentina"

**Table A. GLMM developed for *Ae. aegypti* starting with the variable highV.**

| Model | Variables | AICc |
|---|---|---|
| Ma | highV | 7750.9 |
| Ma1 | highV+lowV | 7752.9 |
| Ma2 | highV+soil | 7745.9 |
| Ma3 | highV+ndvimin | **7683.7** |
| Ma4 | highV+ndwimin | 7749.4 |
| MA5 | highV+ndbimax | |
| Ma6 | highV*lowV | 7706.6 |
| Ma7 | highV*soil | 7687.3 |
| Ma8 | highV*ndvimin | **7665.6** |
| Ma9 | highV*ndwimin | 7748.6 |
| Ma10 | highV*ndbimax | 7726.0 |
| Null | - | 7771.8 |

**Table B. GLMM developed for *Ae. aegypti* starting with the variable soil.**

| Model | Variables | AICc |
|---|---|---|
| Ms | soil | 7753.7 |
| Ms1 | soil+highV | 7745.9 |
| Ms2 | soil+lowV | 7738.2 |
| Ms3 | soil+urban | 7751.6 |
| Ms4 | soil+ndvimin | 7683.6 |
| Ms5 | soil+ndwimax | 7754.9 |
| Ms6 | soil+ndbimin | 7754.5 |
| Ms7 | soil+ndbimax | 7701.5 |
| Ms8 | soil*highV | 7687.3 |
| Ms9 | soil*lowV | 7730.3 |
| Ms10 | soil*urban | 7735.1 |
| Ms11 | soil*ndvimin | **7643.2** |
| Ms12 | soil*ndwimax | 7750.7 |
| Ms13 | soil*ndbimin | 7742.8 |
| Ms14 | soil*ndbimax | 7700.7 |
| Null | - | 7771.8 |

**Table C. GLMM developed for *Ae. aegypti* starting with the variable ndvimin.**

| Model | Variables | AICc |
|---|---|---|
| Mv | ndvimin | 7683.6 |
| Mv1 | ndvimin+highV | 7683.7 |
| Mv2 | ndvimin+lowV | 7682.0 |
| Mv3 | ndvimin+soil | 7683.6 |
| Mv4 | ndvimin+urban | 7678.8 |
| Mv5 | ndvimin +ndwimax | 7685.4 |
| Mv6 | ndvimin +ndbi | 7651.4 |
| Mv7 | ndvimin +ndbimin | 7682.8 |
| Mv8 | ndvimin +ndbimax | 7639.3 |
| Mv9 | ndvimin*highV | 7665.6 |
| Mv10 | ndvimin*lowV | 7672.8 |
| Mv11 | ndvimin*soil | 7643.2 |
| Mv12 | ndvimin*urban | 7678.6 |
| Mv13 | ndvimin*ndwimax | 7678.6 |
| Mv14 | ndvimin*ndbi | 7648.1 |
| Mv15 | ndvimin*ndbimin | 7683.0 |
| Mv16 | ndvimin*ndbimax | **7633.7** |
| Null | - | 7771.8 |

**Table D. GLMM developed for *Ae. aegypti* starting with the variable ndbi.**

| Model | Variables | AICc |
|---|---|---|
| Mb | ndbi | 7764.4 |
| Mb1 | ndbi+lowV | 7766.0 |
| Mb2 | ndbi+ndvimin | 7651.4 |
| Mb3 | ndbi+ndwimax | 7764.9 |
| Mb4 | ndbi*lowV | 7767.1 |
| Mb5 | ndbi*ndvimin | **7648.1** |
| Mb6 | ndbi*ndwimax | 7763.6 |
| Null | - | 7771.8 |

**Table E. GLMM developed for *Ae. aegypti* starting with the variable ndbimax.**

| Model | Variables | AICc |
|---|---|---|
| Mm | ndbimax | 7755.0 |
| Mm1 | ndbimax+highV | 7728.3 |
| Mm2 | ndbimax+lowV | 7755.0 |
| Mm3 | ndbimax+soil | 7701.5 |
| Mm4 | ndbimax+urban | 7756.6 |
| Mm5 | ndbimax+ndvi | 7746.1 |
| Mm6 | ndbimax+ndvimin | 7639.3 |
| Mm7 | ndbimax+ndvimax | 7755.3 |
| Mm8 | ndbimax+ndwi | 7750.9 |
| Mm9 | ndbimax+ndwimin | 7752.4 |
| Mm10 | ndbimax+ndwimax | 7756.9 |
| Mm11 | ndbimax*highV | 7726.0 |
| Mm12 | ndbimax*lowV | 7750.9 |

| | | |
|---|---|---|
| Mm13 | ndbimax*soil | 7700.7 |
| Mm14 | ndbimax*urban | 7757.5 |
| Mm15 | ndbimax*ndvi | 7747.8 |
| Mm16 | ndbimax*ndvimin | **7633.7** |
| Mm17 | ndbimax*ndvimax | 7757.0 |
| Mm18 | ndbimax*ndwi | 7753.0 |
| Mm19 | ndbimax*ndwimin | 7751.8 |
| Mm20 | ndbimax*ndwimax | 7759.0 |
| Null | - | 7771.8 |

**Table F. GLMM selected for *Ae. aegypti.***

| Model | Variables | AICc |
|---|---|---|
| Ma3 | highV+ndvimin | 7683.7 |
| Ms11 | soil*ndvimin | 7643.2 |
| Mv16 | ndvimin*ndbimax | **7633.7** |
| Mb5 | ndbi*ndvimin | 7648.1 |
| Mm16 | ndbimax*ndvimin | **7633.7** |

**Table G. GLMM developed for *Ae. albopictus* starting with the variable soil.**

| Model | Variables | AICc |
|---|---|---|
| Ms | soil | 3442.5 |
| Ms1 | soil+highV | 3443.2 |
| Ms2 | soil+lowV | 3444.5 |
| Ms3 | soil+urban | 3443.2 |
| Ms4 | soil+ndvimin | 3443.2 |
| Ms5 | soil+ndwimax | 3443.3 |
| Ms6 | soil+ndbimin | 3444.5 |
| Ms7 | soil+ndbimax | 3441.0 |
| Ms8 | soil*highV | 3444.2 |
| Ms9 | soil*lowV | **3440.9** |
| Ms10 | soil*urban | 3445.0 |
| Ms11 | soil*ndvimin | 3441.0 |
| Ms12 | soil*ndwimax | 3443.6 |
| Ms13 | soil*ndbimin | 3442.8 |
| Ms14 | soil*ndbimax | **3440.9** |
| Null | - | 3445.4 |

**Fig. A. Photo taken at sampling sites when mosquito larvae were collected.**

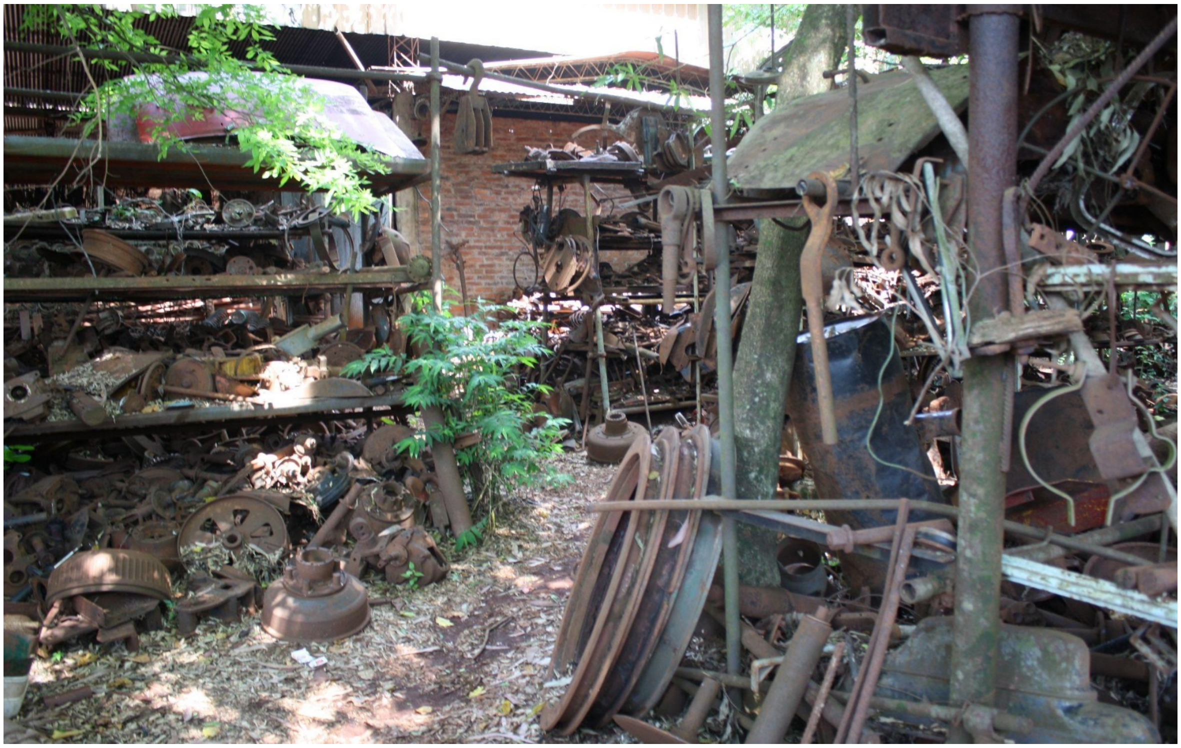
